# DEPICT enables sample-centric therapeutic prioritization across evolving and spatially heterogeneous tumor states

**DOI:** 10.64898/2026.10.07.756157

**Authors:** Shiwen Ma, Shi Yan, Xiaohu Wen, Chenxi Yu, Honghao Ma, Wenzhong Teng, Xuantong Zhou, Jingxi Gao, Siyun Wang, Mingxuan Li, Yanyi Huang, Kui Han, Nan Wu, Bing Liu, Chunhong Zheng

## Abstract

Accurate treatment selection remains a major challenge in precision oncology because tumors with similar genomic alterations can respond differently to therapy. Most drug-response models evaluate individual drugs or drug-sample pairs, whereas clinical decision-making requires therapies to be prioritized within each tumor context. Here we present DEPICT (Drug Efficacy Prediction via Integrated ConText), a sample-centric deep learning framework that reframes drug-response prediction as the relative prioritization of therapies within an individual tumor context. DEPICT integrates transcriptomic and mutational features with prior alteration-therapy relationship and was trained on 231,880 cell line-drug pairs comprising 941 cancer cell lines and 289 compounds. Without retraining, DEPICT retained predictive performance in independently profiled cell lines, experimentally induced drug-resistant states and matched primary lung tumors and patient-derived organoids. Representations learned exclusively from bulk pharmacogenomic data further captured progressive drug-response states at single-cell resolution and outperformed existing single-cell drug-response inference methods. Applied to spatial transcriptomic data, DEPICT resolved clone-associated therapeutic heterogeneity within individual lung tumors, and predicted regional differences were recapitulated in matched multiregional organoid drug-response assays. These findings establish sample-centric therapeutic prioritization as a transferable strategy for modeling dynamic and spatially heterogeneous drug vulnerabilities across biological scales.

## Introduction

Accurate treatment selection remains a central challenge in precision oncology. Genomic profiling has transformed cancer classification and enabled therapies directed against actionable alterations^1-3^, yet genotype alone incompletely explains therapeutic response^4^. Patients harboring the same driver alteration can experience markedly different treatment outcomes^5, 6^, and the activity of cytotoxic and targeted therapies is further shaped by transcriptional state, pathway activity and acquired adaptive programs^7, 8^. Effective treatment prioritization therefore requires models that account not only for static genomic alterations but also for the molecular state in which those alterations occur
.

Computational approaches have increasingly been used to predict anticancer drug response from molecular profiles^9-15^. Early machine-learning models inferred quantitative measures such as half-maximal inhibitory concentration (IC50) and area under the dose-response curve (AUC) from gene-expression or genomic features^16^, whereas more recent deep-learning approaches have enabled joint representation of high-dimensional molecular and pharmacological information^17-22^. However, many approaches are optimized either for individual drugs or for arbitrary drug-sample pairs on a shared absolute response scale^23-25^. Such formulations can be sensitive to differences in experimental platform, dose range and pharmacological measurement and do not directly reflect the comparative nature of treatment selection, in which the clinically relevant question is often which therapy should be prioritized for a particular tumor.

This challenge becomes more pronounced when drug vulnerability is considered at cellular and spatial resolution. Single-cell methods have begun to resolve such heterogeneity through gene-signature scoring, pharmacogenomic integration, transfer learning and related approaches^26-40^, but many require modality-specific modelling or adaptation to sparse single-cell measurements. Spatial transcriptomics further extends this problem by resolving molecularly distinct regions within intact tumors, for which corresponding drug-response measurements are rarely available. Consequently, it remains unclear whether treatment-relevant representations learned from large-scale pharmacogenomic data can transfer without retraining across experimental systems and molecular resolutions while retaining information about evolving and spatially restricted therapeutic vulnerabilities.

Here we developed DEPICT, a sample-centric deep learning framework that directly formulates drug-response prediction as within-sample therapeutic prioritization. DEPICT integrates gene expression, somatic mutation features and established alteration-therapy relationships to learn representations linking tumor state to relative drug vulnerability. We evaluated the framework across independently profiled cell lines, experimentally induced resistance models and primary tumors with matched patient-derived organoids. We further tested whether representations learned exclusively from bulk pharmacogenomic data could be applied directly to single-cell and spatial transcriptomic profiles without model retraining. Across these settings, DEPICT captured treatment-associated state transitions, resolved clone-associated spatial therapeutic heterogeneity and predicted regional differences that were reproduced in matched multiregional organoid assays. These analyses establish a framework for modeling therapeutic vulnerability as a dynamic and heterogeneous property of tumor state across biological scales.

## Results

### DEPICT formulates drug response as sample-centric therapeutic prioritization

We developed DEPICT, a deep learning framework designed to prioritize candidate therapies within an individual molecular context rather than predict drug-sample responses independently (Fig. 1a). The framework comprises a multi-omic sample encoder, a drug encoder, a sample-drug scoring network and a target-bias network. Gene-expression and binary mutation features from 941 Cancer Cell Line Encyclopedia (CCLE) cell lines were projected into a shared latent space through modality-specific multilayer perceptrons (MLPs) and a learnable gating mechanism (Extended Data Fig. 1a, b). Drug representations were learned jointly from pharmacological response measurements for 289 compounds in the Genomics of Drug Sensitivity in Cancer (GDSC) dataset. Established alteration-therapy relationships were incorporated through a target-bias term for molecularly matched drug-sample pairs. In total, DEPICT was trained on 231,880 cell line-drug pairs and generated a ranked therapeutic profile for each sample (Extended Data Fig. 1c, d).

**Fig. 1.**
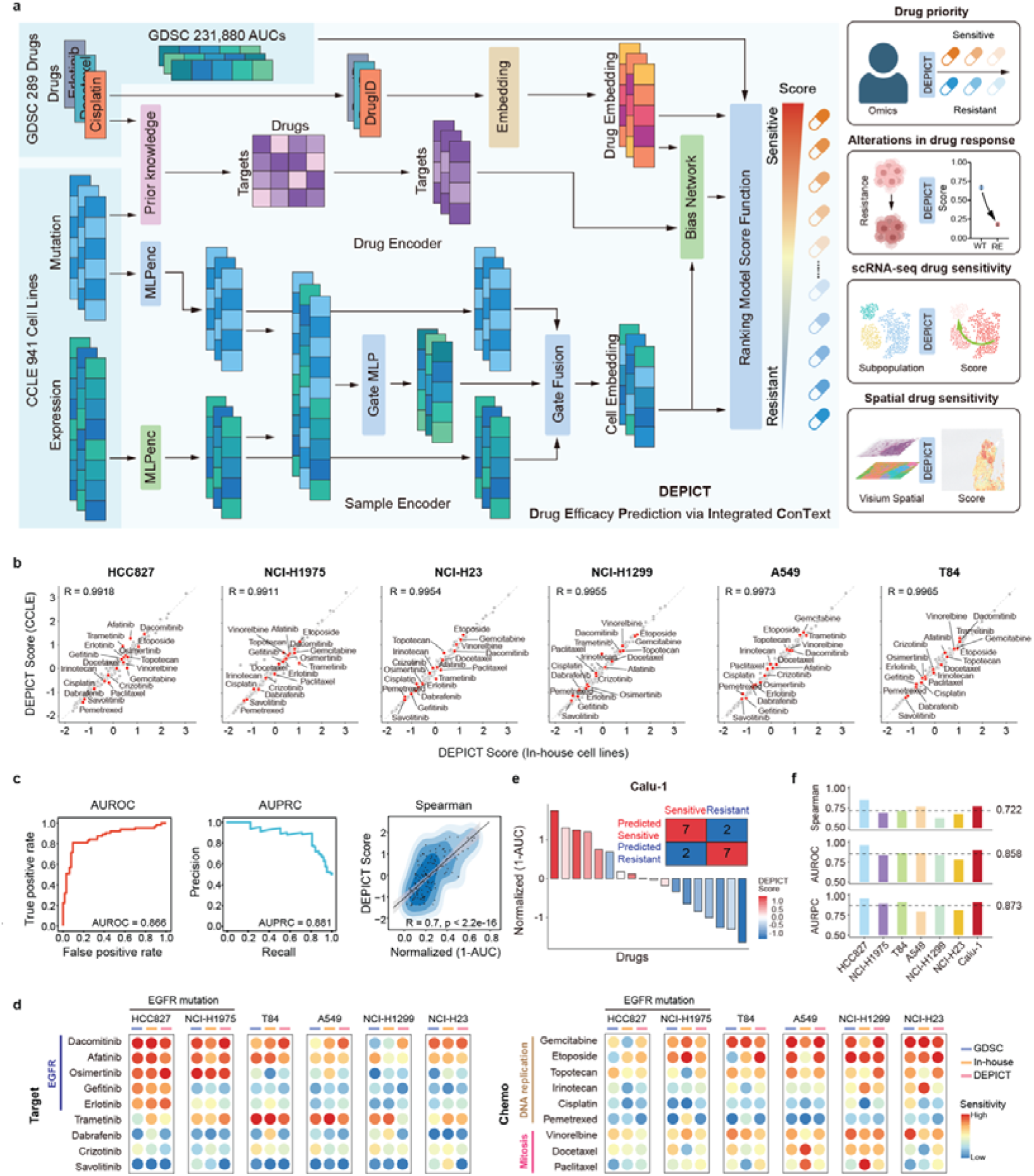
Overview of the DEPICT framework and performance evaluation in lung cancer cell line datasets. **a**, Schematic of the DEPICT. Gene expression and binary mutation features from 941 CCLE cell lines are projected into a shared latent space and fused through a learnable gating mechanism. Drug representations are learned from GDSC response data, and alteration-therapy relationships are incorporated through a target bias network. The model scores candidate drugs within each sample and returns a ranked therapeutic profile. **b**, Correlation of DEPICT-predicted drug sensitivity scores derived from in-house and CCLE multi-omic profiles across six matched lung cancer cell lines. Spearman correlation coefficients are shown. Red points indicate the 18 drugs tested experimentally. **c**, Overall predictive performance across all cell line-drug pairs, assessed by AUROC, AUPRC and Spearman correlation. **d**, Drug sensitivity profiles from GDSC, in-house measurements and DEPICT predictions across six cell lines and 18 drugs. **e**, Predicted and experimentally measured responses in Calu-1, which was excluded from model training. DEPICT correctly classified seven relatively sensitive and seven resistant drugs. **f**, Performance metrics for individual cell lines across the 18 drugs. Dashed lines indicate the corresponding cohort averages.

We next sought to align model optimization with this sample-centric formulation. Conventional objectives based on cross-entropy, mean squared error or binary cross-entropy primarily optimized absolute or categorical response prediction, whereas margin-ranking loss directly modeled the relative ordering of candidate therapies. We therefore developed a composite binary margin-ranking objective that first distinguishes relatively sensitive from resistant drugs across the candidate space and then refines the ordering of drugs within the sensitive subset. All objectives showed stable training and validation dynamics (Extended Data Fig. 1e, f), but differed in their ability to recover therapeutic rankings (Extended Data Fig. 1g, h).

Mean squared error and cross-entropy showed comparatively weak overall performance. Binary cross-entropy effectively discriminated sensitive from resistant drugs but provided limited resolution of within-class ordering, whereas margin-ranking loss improved rank-based performance at the expense of classification accuracy and computational efficiency. The composite objective provided the most balanced performance across Kendall, Spearman and Pearson correlations, concordance index, NDCG@10, Precision@10, AUROC and AUPRC and was therefore used for subsequent analyses. Thus, the optimization strategy of DEPICT directly links molecular context to relative therapeutic vulnerability rather than to isolated drug-response estimates.

### DEPICT retains therapeutic rankings across independently profiled lung cancer cell lines

We first tested whether DEPICT preserved therapeutic rankings when molecular measurements were generated independently of the datasets used for model development. We performed whole-exome and RNA sequencing of six CCLE-matched lung cancer cell lines and compared the resulting profiles with their corresponding public data. Driver-alteration patterns were highly concordant between the independently generated in-house and CCLE datasets (Extended Data Fig. 2a), and transcriptomic profiles showed mean Spearman correlations above 0.8 (Extended Data Fig. 2b). DEPICT scores derived from the independently generated molecular profiles closely recapitulated those obtained from the corresponding CCLE profiles (Fig. 1b), indicating robustness to experimental and dataset-specific variation.

We then measured responses to 18 targeted and cytotoxic agents across the lung cancer cell-line panel (Extended Data Fig. 2c). Independently measured in-house AUC and IC50 values were concordant with GDSC measurements (Extended Data Fig. 2d, e). Using experimental AUC values as ground truth, DEPICT achieved an AUROC of 0.866, an AUPRC of 0.881 and a Spearman correlation of 0.70 between predicted and measured responses (Fig. 1c). The predicted drug-sensitivity profiles further reproduced compound-specific response patterns observed in both GDSC and the independent in-house assays (Fig. 1d). Consistent with established molecular dependencies, EGFR-mutant cell lines showed preferential sensitive to dacomitinib, afatinib, osimertinib, gefitinib and erlotinib, whereas EGFR-wild-type models showed relatively greater sensitivity to gemcitabine and etoposide.

To more stringently assess performance in a molecular context excluded from model training, we examined Calu-1. DEPICT correctly classified 14 of the 18 tested compounds as relatively sensitive or resistant (Fig. 1e and Extended Data Fig. 2c), with a Spearman correlation of 0.771, an AUROC of 0.901 and an AUPRC of 0.909 (Fig. 1f). These values exceeded the corresponding averages across the remaining independently profiled in-house cell lines. Thus, DEPICT preserved within-sample therapeutic rankings across independently generated molecular measurements and retained predictive performance in a molecular context not represented during training.

### DEPICT tracks state-dependent therapeutic vulnerabilities during acquired resistance

We next tested whether therapeutic representations learned from baseline pharmacogenomic data remained informative as tumor cells entered acquired drug-resistant states. Resistant derivatives were generated by prolonged stepwise drug exposure, including osimertinib- and dacomitinib-resistant NCI-H1975 cells and adagrasib-resistant NCI-H23 and Calu-1 cells (Fig. 2a). Each derivative exhibited a several-fold increase in IC50 for the selecting drug relative to its parental counterpart (Fig. 2b and Extended Data Fig. 3a), confirming acquisition of resistance. Whole-exome and bulk RNA sequencing profiles from the parental and resistant states were subsequently analyzed with DEPICT. Predictive performance remained comparable across parental and resistant populations by AUROC, AUPRC and Spearman correlation (Fig. 2c), indicating that the learned representations remained informative in molecular states that emerged after sustained drug exposure.

**Fig. 2.**
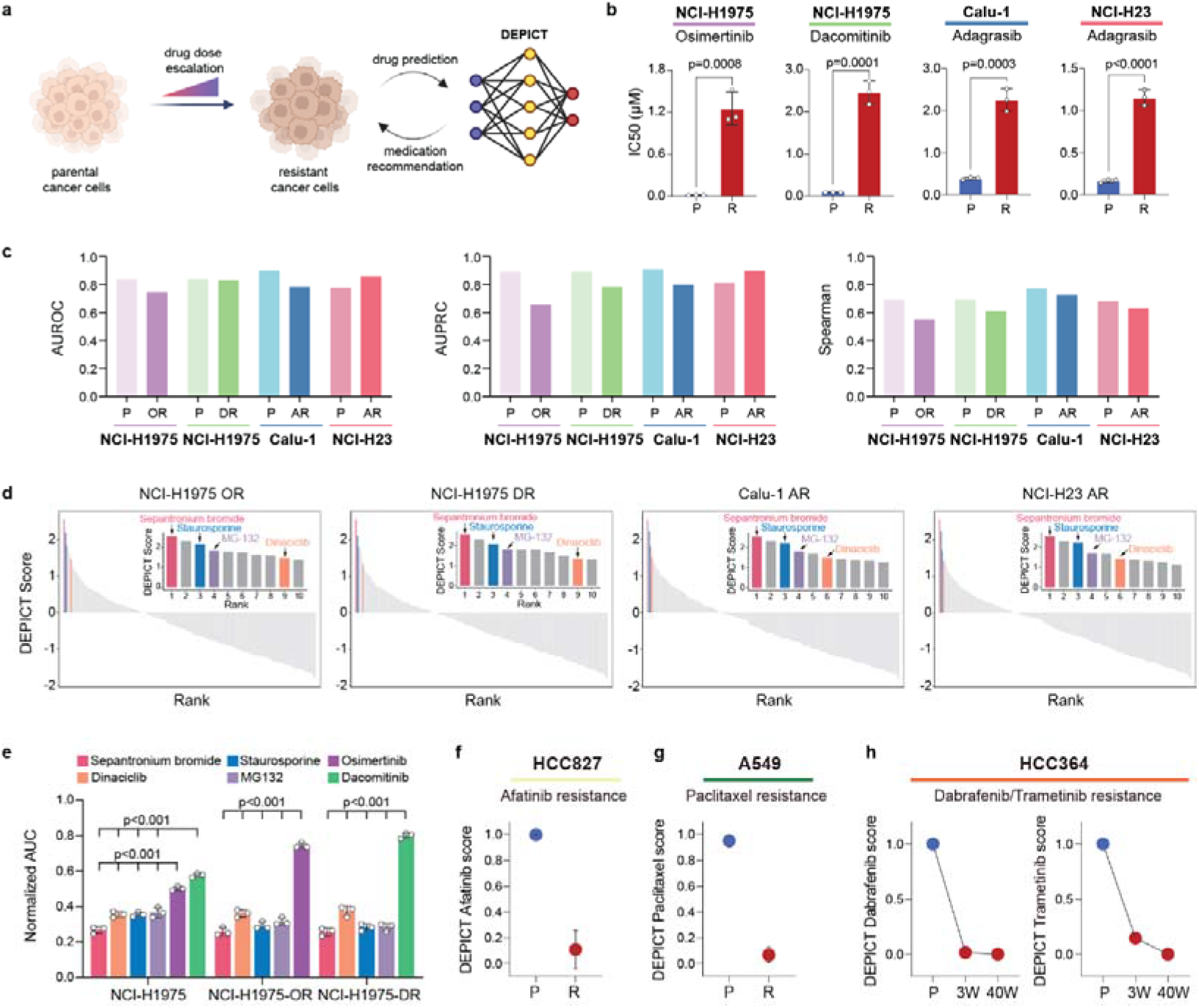
DEPICT captures acquired drug resistance and prioritizes alternative therapeutic vulnerabilities. **a**, Schematic of acquired-resistance model generation by stepwise dose escalation and subsequent DEPICT-based therapeutic prioritization. **b**, IC50 values in parental and resistant NCI-H1975, Calu-1 and NCI-H23 cells. Student’s t-test. **c**, DEPICT performance in parental and resistant states, assessed by AUROC, AUPRC and Spearman correlation. P, parental; OR, osimertinib resistant; DR, dacomitinib resistant; AR, adagrasib resistant. **d**, Ranked DEPICT scores for 289 compounds in the indicated resistant models. Insets show the ten highest-ranked drugs, with recurrent candidates highlighted. **e**, Responses of parental, osimertinib-resistant and dacomitinib-resistant NCI-H1975 cells to selected candidates, shown as normalized AUC. Lower values indicate greater sensitivity. One-way ANOVA. **f**, Predicted afatinib sensitivity in parental and afatinib-resistant HCC827 cells. **g**, Predicted paclitaxel sensitivity in parental and paclitaxel-resistant A549 cells. **h**, Predicted dabrafenib and trametinib sensitivity in HCC364 cells at baseline and after 3 and 40 weeks of resistance induction.

To determine whether DEPICT could identify therapeutic opportunities after resistance acquisition, we ranked all 289 GDSC candidate compounds in each resistant model and recurrently prioritized sepantronium bromide, staurosporine, dinaciclib and MG132 among the highest-scoring candidates (Fig. 2d). Experimental drug-response assays confirmed strong activity of these compounds in the resistant derivatives as well as their parental counterparts, as reflected by low normalized AUC values (Fig. 2e and Extended Data Fig. 3b). Thus, DEPICT not only captured reduced sensitivity to the selecting therapy but also identified pharmacological vulnerabilities retained after resistance acquisition.

We further evaluated independent longitudinal RNA-sequencing datasets using transcriptomic profiles alone. DEPICT predicted a pronounced loss of afatinib sensitivity in HCC827 cells and reduced paclitaxel sensitivity in resistant A549 cells (Fig. 2f, g). In HCC364 cells undergoing prolonged dabrafenib and trametinib exposure, predicted sensitivity to both selecting therapies progressively declined during resistance induction (Fig. 2h). Thus, without retraining on resistant populations, DEPICT tracked treatment-associated shifts in therapeutic vulnerability across experimentally generated and independent longitudinal resistance models.

### Bulk-trained therapeutic representations transfer to patient-derived tumors and organoids

We next examined whether the therapeutic representations learned from cancer cell-line pharmacogenomic data remained informative in patient-derived tumor contexts. Primary lung tumors and matched patient-derived organoids (PDOs) were obtained from 27 patients with non-small cell lung cancer (Fig. 3a and Supplementary Table 1 and 2). PDOs retained major histological and lineage characteristics of their tumors of origin, including expression of thyroid transcription factor 1 (TTF-1) and napsin A in adenocarcinoma and P63 and cytokeratin 5/6 (CK5/6) in squamous cell carcinoma (Fig. 3b and Extended Data Fig. 4a). Whole-exome sequencing further showed preservation of major driver alterations, genome-wide copy number profiles and tumor mutation burden, whereas RNA sequencing revealed high transcriptomic concordance across most matched tumors-PDO pairs (Fig. 3c-f and Extended Data Fig. 4b).

**Fig. 3.**
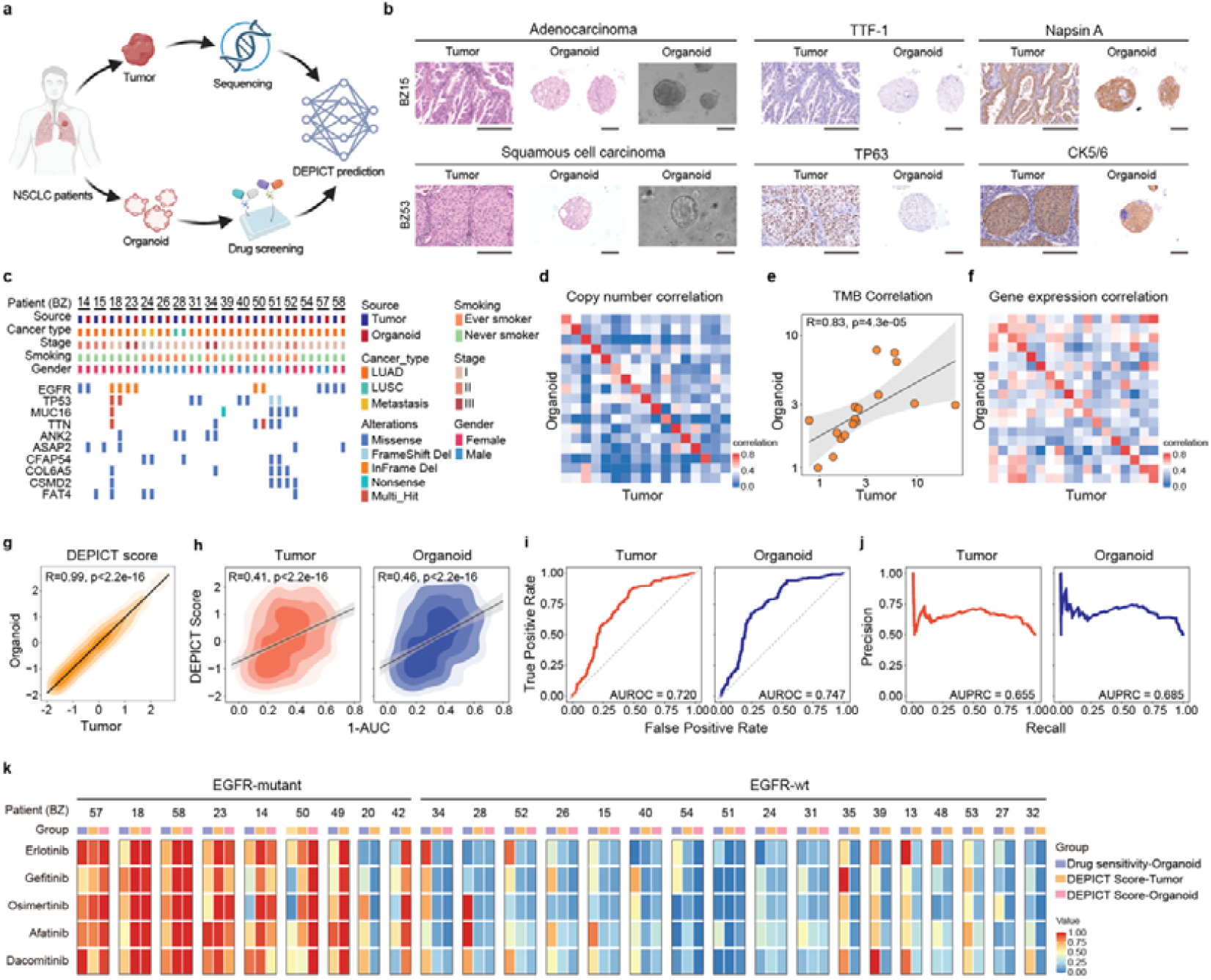
Zero-shot prediction of drug response in patient-derived organoids using DEPICT. **a**, Workflow for collection, multi-omic profiling and DEPICT analysis of matched primary tumors and PDOs from patients with NSCLC. **b**, Representative H&E staining, bright-field imaging and immunohistochemistry showing preservation of histological and lineage features, including TTF-1 and napsin A in adenocarcinoma and TP63 and CK5/6 in squamous cell carcinoma. Scale bars, 100 μm. **c**, Clinical and genomic features of matched tumor-PDO pairs. **d**, Copy number concordance between tumors and PDOs. **e**, Correlation of tumor mutation burden between tumors and PDOs. **f**, Gene-expression concordance between tumors and PDOs. **g**, Correlation of DEPICT scores between matched tumors and PDOs. **h**, Correlation between predicted scores and experimentally measured sensitivity (1 – AUC) using tumor- or PDO-derived inputs. **i**, Receiver operating characteristic curves for tumor- and PDO-based predictions. **j**, Precision-recall curves for tumor- and PDO-based predictions. **k**, Measured PDO sensitivity and DEPICT scores for five EGFR-TKIs, stratified by EGFR mutational status.

Therapeutic profiles generated independently from primary tumors and their matched PDOs were strongly correlated (Fig. 3g), suggesting that DEPICT recovered shared treatment-relevant features across the two experimental contexts. We then screened the PDOs against 18 targeted and chemotherapeutic agents and quantified drug sensitivity as 1-AUC (Extended Data Fig. 4c). DEPICT scores generated from either primary-tumor or PDO molecular profiles correlated significantly with experimentally measured drug sensitivity (Fig. 3h). Tumor-derived predictions achieved an AUROC of 0.720 and an AUPRC of 0.655, whereas PDO-derived predictions achieved an AUROC of 0.747 and an AUPRC of 0.685 (Fig. 3i, j).

Both tumor- and PDO-derived predictions further reproduced genotype-associated pharmacological differences. Samples carrying EGFR mutations showed greater measured and predicted sensitivity to EGFR tyrosine kinase inhibitors than EGFR-wild-type samples (Fig. 3k). These findings indicate that therapeutic representations learned exclusively from cell-line pharmacogenomic data remain informative when applied to primary tumors and patient-derived models without model retraining.

### DEPICT captures resistance-associated states at single-cell resolution

We therefore tested whether therapeutic representations learned exclusively from bulk pharmacogenomic data could capture treatment-associated states at single-cell resolution without explicit transfer learning or single-cell-specific retraining. We first analyzed A375 cells sampled during stepwise dabrafenib exposure from 0 to 4,000 nM^41^. Single-cell transcriptomic profiles showed progressive state transitions across treatment conditions (Fig. 4a, b), accompanied by increased copy-number variation scores at higher drug concentrations (Extended Data Fig. 5a, b). DEPICT recapitulated this progression, with predicted dabrafenib sensitivity progressively decreasing with increasing treatment exposure (Fig. 4c and Extended Data Fig. 5c).

**Fig. 4.**
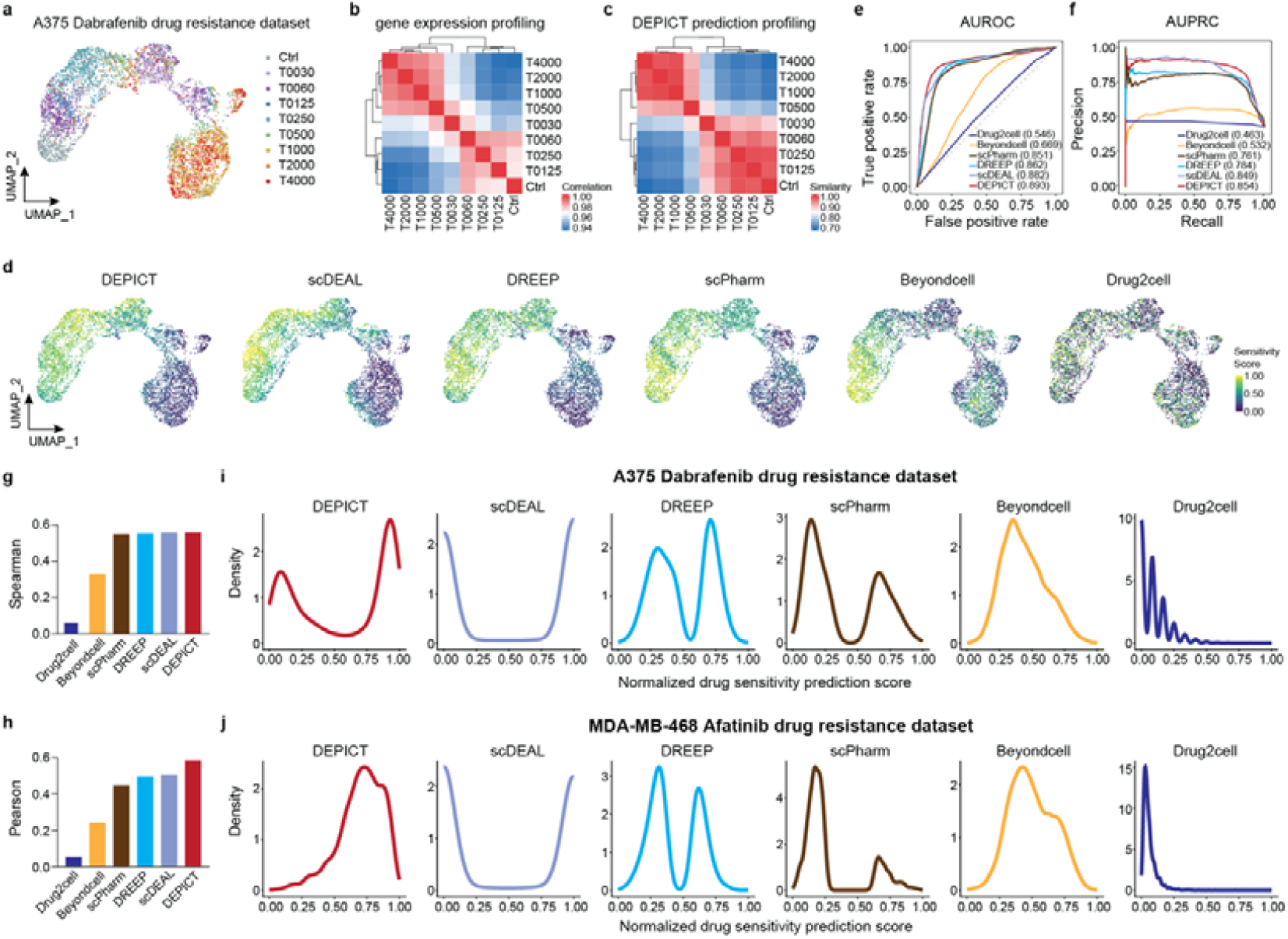
DEPICT resolves drug sensitivity dynamics at single-cell resolution. **a**, UMAP of A375 cells exposed to stepwise dabrafenib escalation. **b**, Correlation heatmap of gene expression profiles across treatment conditions. **c**, Similarity heatmap of DEPICT-predicted drug sensitivity profiles across treatment conditions. **d**, UMAPs showing single-cell sensitivity scores predicted by DEPICT, scDEAL, DREEP, scPharm, Beyondcell and Drug2cell. **e, f**, AUROC (**e**) and AUPRC (**f**) for distinguishing sensitive and resistant cells. **g, h**, Spearman (**g**) and Pearson (**h**) correlations between predicted scores and treatment-associated sensitivity states. **i, j**, Density distributions of normalized prediction scores generated by the indicated methods in the A375 dabrafenib (**i**) and MDA-MB-468 afatinib (**j**) resistance dataset.

We benchmarked DEPICT against five methods used for single-cell drug-response inference, including scDEAL, DREEP, scPharm, Beyondcell and Drug2cell. DEPICT achieved the highest discrimination performance in the A375 resistance series, with an AUROC of 0.893 and an AUPRC of 0.854 (Fig. 4d-f). It also showed the strongest Spearman and Pearson correlations between predicted scores and treatment-associated sensitivity states (Fig. 4g, h), indicating recovery of the progressive resistance trajectory rather than only separation of the exposure endpoints.

We next evaluated an independent MDA-MB-468 single-cell dataset of stepwise afatinib resistance induction^42^ (Extended Data Fig. 5d). The initial populations contained coexisting sensitive and resistant cells, creating a setting in which the two states were not readily separable by global transcriptomic structure alone. DEPICT again achieved the highest AUROC and AUPRC among the methods tested (Extended Data Fig. 5e, f). The distributions of predicted sensitivity scores also differed substantially among methods. Drug2cell and Beyondcell produced predominantly unimodal distributions, scPharm and DREEP generated pronounced bimodal patterns and scDEAL yielded near-binary predictions. By contrast, DEPICT produced a bimodal distribution in the A375 model but a predominantly unimodal distribution in MDA-MB-468, reflecting the distinct state structure of the two datasets (Fig. 4i, j and Extended Data Fig. 5g, h). Together, these results show that DEPICT can recover the continuous and context-dependent transcriptional states associated with drug sensitivity and resistance at single-cell resolution.

### DEPICT resolves and functionally validates spatial therapeutic heterogeneity

We next asked whether treatment-associated molecular states were spatially organized within intact tumors. DEPICT was applied to Visium spatial transcriptomic profiles from three lung adenocarcinomas^43^. Tumor-associated markers delineated malignant regions, and fastCNV analysis revealed substantial intratumoral copy-number heterogeneity (Extended Data Fig. 6a), enabling malignant regions to be partitioned into spatially organized clones.

In case 1, two major clones were identified (Fig. 5a-c). Clone 2 showed chromosome 7 gain and chromosome 9 loss (Fig. 5d), encompassing *EGFR* and *CDKN2A*, respectively, and these genomic differences were accompanied by increased *EGFR* and reduced *CDKN2A* expression (Fig. 5e, f). DEPICT predicted substantially greater sensitivity of clone 2 to five EGFR tyrosine kinase inhibitors, including afatinib, dacomitinib, gefitinib, erlotinib and osimertinib, than clone 1 (Fig. 5g, h), consistent with an EGFR-dependent therapeutic state^44-46^.

**Fig. 5.**
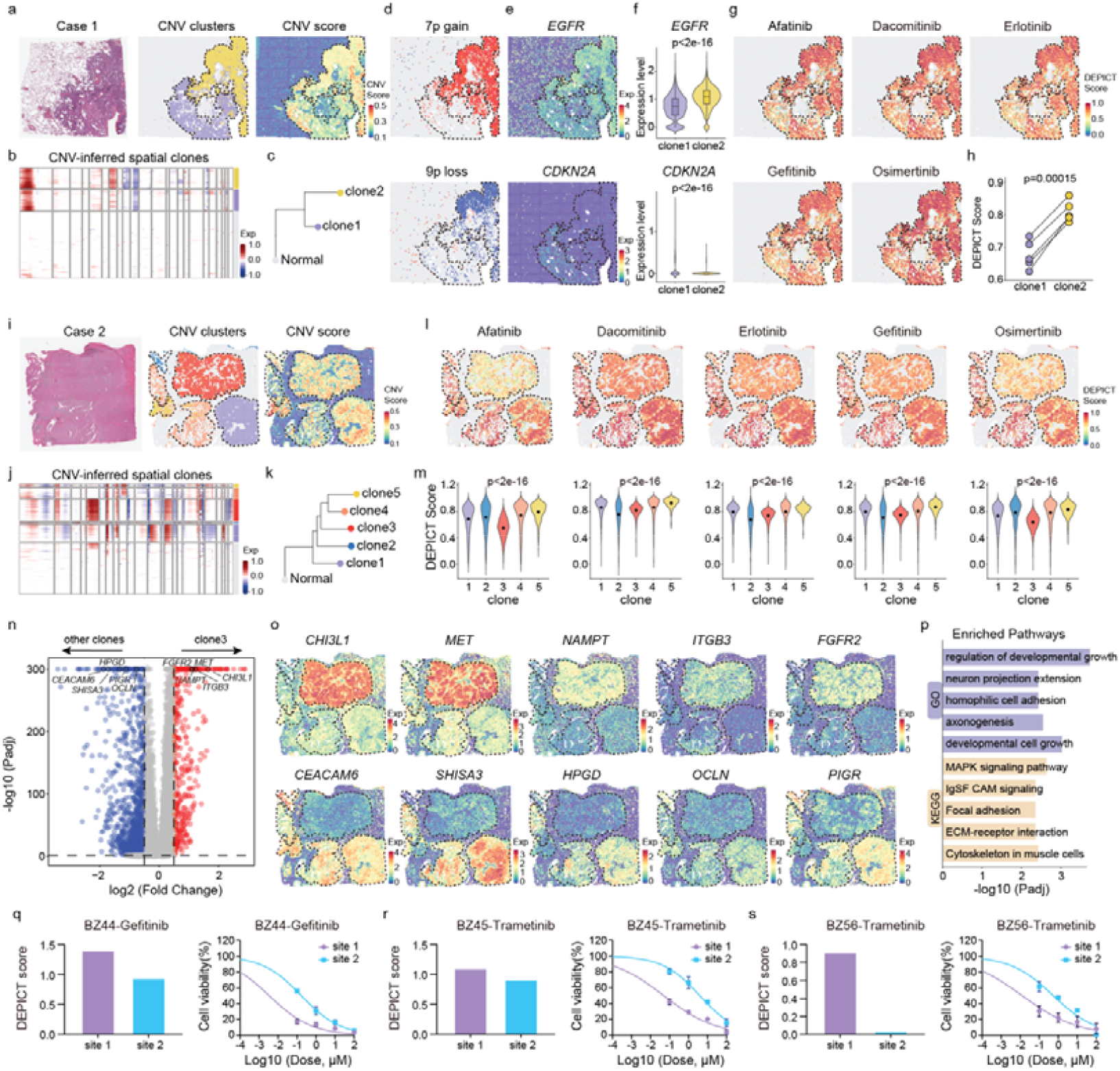
DEPICT resolves spatially heterogeneous drug sensitivity across tumor clones in lung adenocarcinoma. **a**, H&E staining, fastCNV-inferred clone assignment and CNV score for case 1, identifying two spatial clones. **b**, fastCNV-inferred CNV heatmap for case 1. **c**, Phylogenetic tree of normal regions and tumor clones in case 1. **d**, Spatial distribution of chromosome 7p gain and chromosome 9p loss. **e**, Spatial expression of EGFR and CDKN2A. **f**, EGFR and CDKN2A expression in the two clones. Two-sided Wilcoxon rank-sum tests. **g**, Predicted sensitivity maps for afatinib, dacomitinib, erlotinib, gefitinib and osimertinib. **h**, Mean EGFR-TKI sensitivity scores in the two clones. Two-sided paired Student’s t-test. **i**, H&E staining, clone assignment and CNV score for case 2, identifying five spatial clones. **j**, fastCNV-inferred CNV heatmap for case 2. **k**, Phylogenetic tree of normal regions and tumor clones in case 2. **l**, Predicted sensitivity maps for five EGFR-TKIs. **m**, EGFR-TKI sensitivity scores across clones. Two-sided Kruskal-Wallis test. **n**, Differentially expressed genes between clone 3 and the remaining clones (|log2 fold change| > 1; adjusted P < 0.05). **o**, Spatial expression of representative clone3-associated genes. **p**, Gene Ontology and KEGG enrichment of clone 3-associated genes. **q**, DEPICT-predicted gefitinib sensitivity in two spatially distinct regions of BZ44, together with matched organoid dose-response curves validating region-specific differences in gefitinib sensitivity. **r**, DEPICT-predicted trametinib sensitivity in two regions of BZ45, with matched organoid dose-response assays confirming differential trametinib responses between sites. **s**, DEPICT-predicted trametinib sensitivity in two regions of BZ56, with matched organoid dose-response assays validating pronounced regional differences in trametinib sensitivity.

Case 2 contained five spatially distinct malignant clones (Fig. 5i-k), one of which, clone 3, exhibited markedly reduced predicted sensitivity to EGFR-targeted therapies, most prominently afatinib (Fig. 5l, m). Compared with the remaining tumor clones, clone 3 showed increased expression of *CHI3L1, FGFR2, MET, ITGB3* and *NAMPT* and reduced expression of *CEACAM6, SHISA3, HPGD, PIGR* and *OCLN* (Fig. 5n, o), which are reported to be associated with therapeutic resistance^47-52^. Gene Ontology and KEGG analyses further revealed enrichment of programs related to cell growth, MAPK-signaling, focal adhesion and extracellular matrix-receptor interactions (Fig. 5p), consistent with a spatially restricted state characterized by bypass signaling, extracellular matrix remodeling and reduced EGFR-TKI sensitivity.

In case 3, two spatially adjacent clones exhibited reciprocal predicted vulnerabilities to cytotoxic chemotherapy and EGFR-targeted therapy (Extended Data Fig. 6b-d). Clone 1 showed higher predicted sensitivity to multiple chemotherapeutic agents but lower sensitivity to EGFR-TKIs, whereas clone 2 exhibited the opposite pattern (Extended Data Fig. 6e-g). Differential expression analysis revealed that clone 1 was characterized by increased expression of *TP63, CD24, AREG, EREG* and *ROS1*, whereas clone 2 showed increased expression of *SERPINA3, IGFBP5, TMPRSS4, CCND1* and *SMAD3* (Extended Data Fig. 6h, i), alterations that have been associated with resistance to corresponding targeted or cytotoxic therapies^53-58^. Gene set enrichment analysis further identified an EGFR-TKI-tolerant program in clone 1 and a doxorubicin-resistance program in clone 2 (Extended Data Fig. 6j), supporting spatially neighboring tumor populations can occupy divergent therapeutic states despite arising within the same lesion.

To functionally validate the region-specific differences predicted from spatial transcriptomic data, we collected spatially distinct tumor regions from three additional cases and independently profiled each region by RNA sequencing and whole-exome sequencing. Matched patient-derived organoids were established from the corresponding regions and subjected to drug sensitivity testing. DEPICT predicted marked regional differences in response to selected therapies in all three cases, including gefitinib in BZ44 and trametinib in BZ45 and BZ56. Importantly, matched organoid drug-response assays reproduced these region-specific differences (Fig. 5q-s), providing functional evidence that spatially distinct regions within the same tumor can harbor divergent therapeutic sensitivities. Together with the single-cell and spatial analyses, the multiregional validation establishes therapeutic response as a dynamic and spatially heterogeneous property of tumor state that can be modeled using representations learned from large-scale pharmacogenomic data.

## Discussion

In this study, we developed DEPICT, a sample-centric framework that prioritizes anticancer therapies within an individual molecular context. Rather than treating each drug-sample pair as an independent prediction task, DEPICT evaluates candidate therapies relative to one another for the same molecular profile. By integrating transcriptomic and mutational features with established alteration-therapy relationships, DEPICT learns therapeutic rankings that can be applied across molecular contexts with different data availability.

A central finding of this study is that treatment-relevant representations learned from large-scale bulk pharmacogenomic data remained informative in molecular states not explicitly represented during training. DEPICT retained performance in independently profiled cell lines and patient-derived organoids and captured changes associated with experimentally induced drug resistance. In longitudinal resistance models, predicted sensitivity decreased progressively as cells adapted to the selecting therapy, indicating that the framework did not simply distinguish fixed sensitive and resistant classes but tracked treatment-associated changes along evolving molecular states. These observations support the view that therapeutic vulnerability is a dynamic property of tumor state rather than a static attribute determined solely by genotype.

The extension to single-cell data further showed that this therapeutic-state representation could operate at cellular resolution without explicit single-cell model training or domain adaptation. Across two independent resistance models^41^, DEPICT reconstructed context-dependent drug-response states and compared favorably with methods specifically developed for single-cell drug-response inference. We nevertheless interpret these analyses as evidence that DEPICT captures transcriptional states associated with therapeutic sensitivity and resistance, rather than as direct measurements of the pharmacological response of every individual cell. Functional measurements coupled to single-cell profiling will be required to define the limits of this resolution more precisely.

The spatial analyses provide a complementary view of intratumoral therapeutic heterogeneity. Genetically and transcriptionally distinct tumor clones within the same lesion were associated with divergent predicted responses to targeted therapy and chemotherapy. These differences were not restricted to canonical driver alterations but were accompanied by bypass-signaling, metabolic and extracellular-matrix programs associated with resistant tumor states^47-52^. Importantly, independent multiregional sampling and matched organoid assays demonstrated that molecularly distinct regions from the same tumor could exhibit corresponding functional differences in drug response. These experiments move beyond spatial visualization of predicted sensitivity and provide direct evidence that regional tumor states can encode distinct therapeutic vulnerabilities. Such heterogeneity offers a potential explanation for incomplete responses to uniform treatment^6, 59^ and highlights the importance of considering spatial tumor architecture when defining therapeutic strategies.

Several limitations remain. DEPICT was trained predominantly on in vitro cancer cell-line pharmacogenomic data and therefore cannot capture all influences of the tumor microenvironment, pharmacokinetics or host physiology on treatment response^60, 61^. The patient-derived cohort was modest, and the present study did not directly test whether pretreatment DEPICT rankings predict clinical response or survival in patients receiving the corresponding therapies. Moreover, because drugs are represented by learned embeddings, the current implementation prioritizes compounds represented during model training rather than providing zero-shot prediction for entirely unseen drugs. Finally, functional validation at single-cell and high-resolution spatial scales remains limited.

Together, our findings establish sample-centric therapeutic prioritization as a framework for modeling drug vulnerability across evolving and spatially heterogeneous tumor states. Rather than treating therapeutic response as a fixed property of a drug-sample pair, DEPICT captures relative vulnerabilities that persist across experimental systems and can vary among cellular states and spatial tumor regions. This framework provides a basis for connecting pharmacogenomic learning with dynamic and spatial representations of tumor biology and, with further clinical validation, may support more context-aware therapeutic prioritization in precision oncology.

## Methods

### Patient samples and ethics

Thirty patients with non-small cell lung cancer (NSCLC) were enrolled at Peking University Cancer Hospital, including 29 patients with primary lung tumors and 1 patient with pulmonary metastatic disease. Matched tumor-PDO samples from all 30 patients were included in the zero-shot prediction analysis, including three multi-regional tumor-PDO pairs. All participants provided written informed consent. The study was approved by the Ethics Committee of Peking University Cancer Hospital (approval no. 2024KT153) and was conducted in accordance with relevant ethical regulations. Clinical characteristics are provided in Supplementary Tables 1 and 2.

### DEPICT model implementation

DEPICT uses a dual-tower architecture inspired by recommendation and retrieval systems, with task-specific extensions for drug sensitivity prediction. The model comprises a multi-omic sample encoder, a drug encoder, a pairwise scoring network and a target bias network. Gene expression, mutation and drug features are projected into a shared latent space to learn sample-drug representations for response prediction and within-sample ranking.

#### (a) Multi-omic cell encoder

Each sample was represented by a gene expression vector *x*_*exp*_ *∈ R*^*M*^ and a binary mutation vector *x*_*mut*_ *∈ {0,1}*^*K*^ . Independent multilayer perceptrons (MLPs) mapped the two modalities into a shared latent space of dimension *d*:

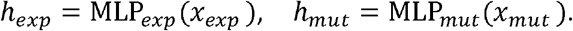

To account for unequal contributions from the two modalities, a learnable gating vector *g*_*θ*_ was computed from the concatenated expression and mutation embeddings:

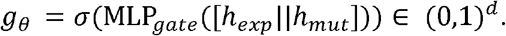

To stabilize fusion and reduce modality collapse, a residual term *r = h*_*exp*_ +*h*_*mut*_ was added. The final sample embedding was defined as follows.

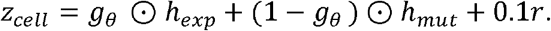

This formulation adaptively balances modality-specific information while retaining complementary signals from gene expression and mutation features.

#### (b) Drug encoder

All drugs were represented by a trainable embedding matrix. Let *N*_*drug*_ denote the number of drugs and 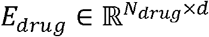 the drug embedding matrix. Drug *j* was mapped into the same latent space as the sample representation:

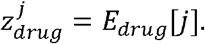

This formulation does not require predefined chemical fingerprints or molecular descriptors and allows drug representations to be learned directly from the training data.

#### (c) Pairwise scoring and bias modeling

For each sample-drug pair *(i, j)*, DEPICT constructed a joint feature vector comprising the sample embedding, the drug embedding, and their element-wise interaction:

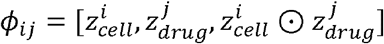

A multilayer perceptron was then used to compute the primary prediction score,

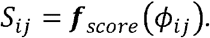

and an auxiliary bias network was used to model mutation-matched targeted therapies, which were relatively sparse in the training data.

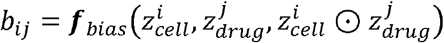

The final prediction score was defined as follows.

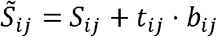

Here, *t*_*ij*_ *∈ {0,1}* indicates whether drug *j* is a known targeted therapy for sample *i*. The bias term was therefore applied only to targeted pairs, whereas non-targeted pairs retained the original score *S*_*ij*_.

#### (d) Composite loss for classification and ranking

DEPICT was optimized using a composite objective that combines binary cross-entropy with margin ranking loss to model both sensitivity classification and within-sample drug prioritization. The binary classification term was defined as follows.

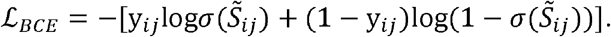

Here, *y*_*ij*_ *∈ {0,1}* denotes the ground-truth sensitivity label. To model the relative drug efficacy within a sample, drugs *j* and *k* were ordered according to their observed 1-AUC values. When drug j was more sensitive than drug k in sample *i*, the model was optimized to separate their predicted scores by at least margin m.

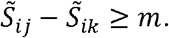

The corresponding margin ranking loss was defined as follows.

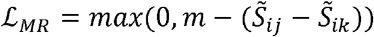

Drug pairs were generated by positive-negative sampling, and the total objective combined the classification and ranking terms.

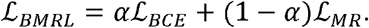

The coefficient *α∈* [0,1] controls the relative contribution of classification and ranking.

During inference, DEPICT first computes the sample representation from the available multi-omic features and then calculates a score for every candidate drug.

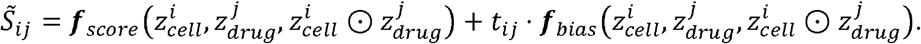

Candidate drugs were ranked by the final score 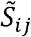 with higher scores indicating greater predicted sensitivity.

### Hyperparameter settings

All experiments used a common hyperparameter configuration. The random seed was fixed at 42, and performance was evaluated using a five-fold cell-line-disjoint cross-validation. Models were trained with the Adam optimizer using a learning rate of 5 × 10^−4^, a batch size of 32 and weight decay of 1 × 10^−4^ for up to 200 epochs. Early stopping was applied after 10 epochs without improvement, with a minimum metric improvement threshold of *δ* = 0.001. Ground-truth AUC values were transformed to 1-AUC, and sample-drug pairs above the 80th percentile were labeled relatively sensitive. Positive and negative examples were balanced by sampling during training.

The composite objective used *α* = 0.5 and a ranking margin of *m* = 0.5. Ranking metrics were evaluated at *K* = 10. The multi-layer perceptrons had hidden dimensions of 512 and 256 and an output embedding dimension of *d= 128*. ReLU activation and dropout of 0.5 were applied to fully connected layers.

### Training datasets

DEPICT was trained using multi-omic profiles from the Cancer Cell Line Encyclopedia (CCLE)^62^ and drug response measurements from the Genomics of Drug Sensitivity in Cancer (GDSC)^16^. Cell lines tested against more than 100 compounds were retained to ensure adequate pharmacological coverage. The final training set comprised 941 cancer cell lines, 289 compounds and 231,880 cell line-drug pairs.

For transcriptomic input, the coefficient of variation was calculated for each gene across retained cell lines, and genes with a coefficient of variation greater than 0.5 were selected. This yielded 3,393 expression features. For genomic input, 611 curated cancer driver mutations from Cell Model Passports were encoded as binary mutation features. Depending on data availability, DEPICT was run using gene expression alone or using integrated expression and mutation features.

### Validation datasets

DEPICT was evaluated in three categories of independent datasets spanning bulk RNA sequencing, single-cell RNA sequencing and Visium spatial transcriptomics. Longitudinal bulk datasets from the Gene Expression Omnibus contained matched parental and drug-resistant cell states and were used to assess resistance-associated transcriptional changes. Drug-treated single-cell datasets were used to test direct inference at cellular resolution without additional model adaptation. Visium datasets were used to evaluate relative drug sensitivity across spatially defined tumor clones. Because matched functional drug-response labels were not available for the spatial datasets, predictions were interpreted alongside copy number, transcriptional and pathway-level evidence.

### Evaluation metrics

Model performance was assessed using Kendall rank correlation, Spearman rank correlation, Pearson correlation, concordance index, area under the receiver operating characteristic curve (AUROC), area under the precision–recall curve (AUPRC), normalized discounted cumulative gain at rank k (NDCG@k) and Precision@k. Kendall rank, Spearman rank and Pearson correlation coefficient were used to quantify rank-based and linear agreement between predicted and observed drug responses. CI was used to assess pairwise ranking accuracy. AUROC and AUPRC were used to evaluate binary discrimination performance. NDCG@k and Precision@k were used to assess the quality of top-ranked predictions. NDCG@10 and Precision@10 were used unless otherwise specified.

### Sample collection and organoid generation

Fresh tumor tissues were transported on ice in Tissue Storage Solution (Miltenyi Biotec). Samples were minced into 1-3 mm^3^ fragments and digested in RPMI-1640 containing collagenase IV, hyaluronidase and DNase I (Thermo Fisher Scientific) at 37 °C for 30 min. The suspension was passed through a 100-μm cell strainer and centrifuged at 300 g for 5 min. Cell pellets were seeded into 24-well plates and cultured with NSCLC organoid culture medium (K2 Oncology). Medium was replaced every 2-3 days, and organoids were passaged every 1-2 weeks.

### Cell culture

HCC827, NCI-H1975, A549, NCI-H1299, NCI-H23, T84 and Calu-1 cells were obtained from Pricella and authenticated by short tandem repeat profiling. Osimertinib-resistant NCI-H1975-OR, dacomitinib-resistant NCI-H1975-DR and adagrasib-resistant Calu-1-AR and NCI-H23-AR derivatives were generated by continuous drug exposure with stepwise dose escalation. Cells were maintained in RPMI-1640 (Pricella) supplemented with 10% Fetal bovine serum (Pricella) and 1% penicillin-streptomycin (Gibco) at 37 ⍰ °C in 5% CO_2_ and were passaged every 2-3 days.

### Compounds

The following compounds were purchased from MedChemExpress (U.S.A), including Gefitinib (HY-50895), Erlotinib (HY-50896), Afatinib (HY-10261), Osimertinib (HY-15772), Dacomitinib (HY-13272), Crizotinib (HY-50878), Cisplatin (HY-17394), Paclitaxel (HY-B0015), Docetaxel (HY-B0011), Pemetrexed (HY-10820), Gemcitabine (HY-17026), Vinorelbine (HY-12053A), Etoposide (HY-13629), lrinotecan (HY-16562), Topotecan (HY-13768A), Savolitinib (HY-15959), Trametinib (HY-10999), Dabrafenib (HY-14660), Adagrasib (HY-130149), Sepantronium bromide (HY-10194), Staurosporine (HY-15141), MG-132 (HY-13259) and Dinaciclib (HY-10492).

### Drug screening

PDOs were dissociated into small spheroids smaller than 70 μm using TrypLE (Gibco) and seeded in triplicate into ultralow-attachment 384-well plates. After 48 h, organoids were exposed for 5 days to drugs concentrations of 0, 0.1, 1, 10 and 100 μM. For cell line assays, 5,000 cells per well were seeded in triplicate into 96-well plates for 24⍰ h and then treated for 48 h with 0, 0.01, 0.1, 1, 10 or 100 μM drug. Viability was quantified using CellTiter-Glo 3D reagent (Promega) according to the manufacturer’s instructions.

### H&E staining and immunohistochemistry

Paraffin-embedded tissue and organoid sections were deparaffinized and rehydrated before H&E staining or immunohistochemistry. For immunohistochemistry, endogenous peroxidase activity was quenched with 3% H_2_O_2_ for 10 min. Antigen retrieval was performed by microwave heating for 15 min in sodium citrate buffer (pH 6.0) or 1 mM EDTA buffer. Sections were washed three times with PBS, blocked with goat serum for 1 h at room temperature and incubated with primary antibodies overnight at 4 ⍰ °C. After washing, sections were incubated with secondary antibodies for 20 min at room temperature, and signals were developed with diaminobenzidine. The primary antibodies applied, including P63 Polyclonal antibody (1:500, Proteintech, 12143-1-AP), Anti-Cytokeratin 5 Monoclonal antibody (EP1601Y) (1:200, abcam, ab52635), TTF-1 Monoclonal antibody (1:100, CST, #23192) and Napsin A Polyclonal antibody (1:200, CST, #62434).

### Whole-exome sequencing and data processing

Genomic DNA was extracted from parental cell line, drug-resistant derivative, PDOs, tumors and matched normal tissues using DNA and RNA Isolation Kits (Genstone Biotech). Matched normal tissue from the same patient served as the germline control. Exonic regions were captured using the iGeneTech T600 V1 probe set, and libraries were sequenced on an Illumina platform to generate 150-bp paired-end reads.

Reads were trimmed and quality filtered using fastp (v0.24.0) and aligned to GRCh38 using BWA-MEM2 (v2.2.1). Alignments were sorted and indexed with Samtools (v1.21). PCR duplicates were marked using MarkDuplicates in GATK (v4.6.0.0), followed by base-quality score recalibration with BaseRecalibrator and ApplyBQSR. Somatic variants were called with GATK Mutect2 with matched normal controls, the gnomAD allele-frequency resource and the 1000 Genomes hg38 panel of normals. Variants were annotated with ANNOVAR. High-confidence somatic variants were retained when the alternate read count was at least 3, normal depth was at least 10 and variant allele fraction was at least 0.05. Copy-number alterations were inferred using CNVkit to generate segmented copy number profiles.

### Bulk RNA sequencing and data processing

Total RNA was isolated from parental cell lines, resistant derivatives, PDOs and tissue specimens using the DNA and RNA Isolation Kit (Genstone Biotech). Poly(A)+ RNA was enriched using poly(T)-oligo-conjugated magnetic beads and used for library preparation. Libraries were sequenced on an Illumina NovaSeq X Plus platform to generate 150-bp paired-end reads.

Reads were processed with fastp (v0.24.0) and aligned to GRCh38 using HISAT2 (v2.2.1). Gene-level counts were obtained with featureCounts from the Subread package using GENCODE release 47 annotations. Expression was normalized to transcripts per million, and log_2_(TPM + 1) values were used for downstream analyses.

### DEPICT drug sensitivity prediction

DEPICT was applied to bulk RNA-seq, single-cell RNA-seq and Visium spatial transcriptomic datasets. Samples with paired expression and mutation data were analyzed in --mode all using log2(TPM+1)-transformed expression values and binary mutation features. Datasets without matched mutation profiles were analyzed in --mode exp using expression alone. Single-cell and spatial expression matrices were log-normalized with Seurat NormalizeData before inference, and predicted scores were added to Seurat metadata. DREEP, scPharm, Beyondcell, Drug2cell and scDEAL were run using their recommended default parameters. For cross-method comparison, scores were rescaled so that higher values indicated greater predicted sensitivity.

### Single-cell and spatial transcriptomic analysis

Single-cell and Visium data were processed in Seurat (v5.4.0) using NormalizeData, FindVariableFeatures, ScaleData, RunPCA, FindNeighbors, FindClusters and RunUMAP. Clustering resolution was selected with clustree. For Visium data, genome-wide copy number profiles were inferred using fastCNV, with normal, immune and stromal regions used as reference populations. Default parameters were used unless otherwise specified.

### Statistical analysis and reproducibility

Statistical analyses were performed using R (v4.4.1) and GraphPad Prism (v9.5.1). Data are presented as mean ± s.d. unless otherwise indicated. Two-group comparisons used unpaired two-tailed Student’s t-test or Wilcoxon rank-sum tests, as appropriate. Comparisons among three or more groups used one-way ANOVA with Dunnett’s multiple-comparisons test or the Kruskal-Wallis test. Pearson and Spearman correlation coefficients assessed linear and monotonic associations, respectively. Multiple testing was controlled using the Benjamini-Hochberg method. Statistical significance was defined as P < 0.05 or FDR q < 0.05. Values below the numerical limit reported by R were presented as P < 2.2 × 10^−16^.

## Data availability

In-house RNA-seq and whole-exome sequencing (WES) data from cell lines, resistant derivatives, tumors and PDOs have been deposited in National Genomics Data Center, under accession number PRJCA069857. External bulk RNA-seq datasets are available under accession numbers GSE304051^52^, GSE277007^53^ and GSE271247^54^. Single-cell datasets are available under GSE247684^55^ and GSE228154^56^. The Visium lung cancer spatial transcriptomics dataset is available under GSE307534^57^. Source data are provided with this paper. Additional information required to reanalyse the data is available from the corresponding authors upon reasonable request.

## Code availability

The source code for DEPICT is available at https://github.com/zhenglab-PKU/DEPICT. The training data, data generated in this study, and processed external validation data have been deposited on Figshare at https://doi.org/10.6084/m9.figshare.33093000. The trained model weights used to generate the results presented in this article are available on Figshare at https://doi.org/10.6084/m9.figshare.33058595.

## Acknowledgements

This work was supported by grants from the Noncommunicable Chronic Diseases-National Science and Technology Major Project (2023ZD0520400 to C.Z. and K.H., and 2023ZD0501700 to N.W.), National Key R&D Program of China (No. 2025YFF1500200 to C.Z.), the National Natural Science Foundation of China (32571101 to C.Z. and 82373082 to N.W.), the Capital’s funds for health improvement and research (No.2026-4-1097 to B.L.), and the Science Foundation of Peking University Cancer Hospital (ZY202604 to S.M. and BJCH2026BJ04 to B.L.). This study was supported by Changping Laboratory.

## Author contributions

S.M., X.W., and C.Z. conceived and designed the study. S.Y., N.W. and B.L. provided study materials. S.M., S.Y., X.W., C.Y., H.M., W.T., X.Z., J.G., S.W., M.L., Y.H. and K.H. performed experiments. S.M., X.W. and C.Z. analyzed and interpreted the data. S.M., S.Y., X.W., N.W., B.L. and C.Z. wrote and revised the manuscript. All authors reviewed and approved the final manuscript.

---

**Extended Data Fig. 1.**
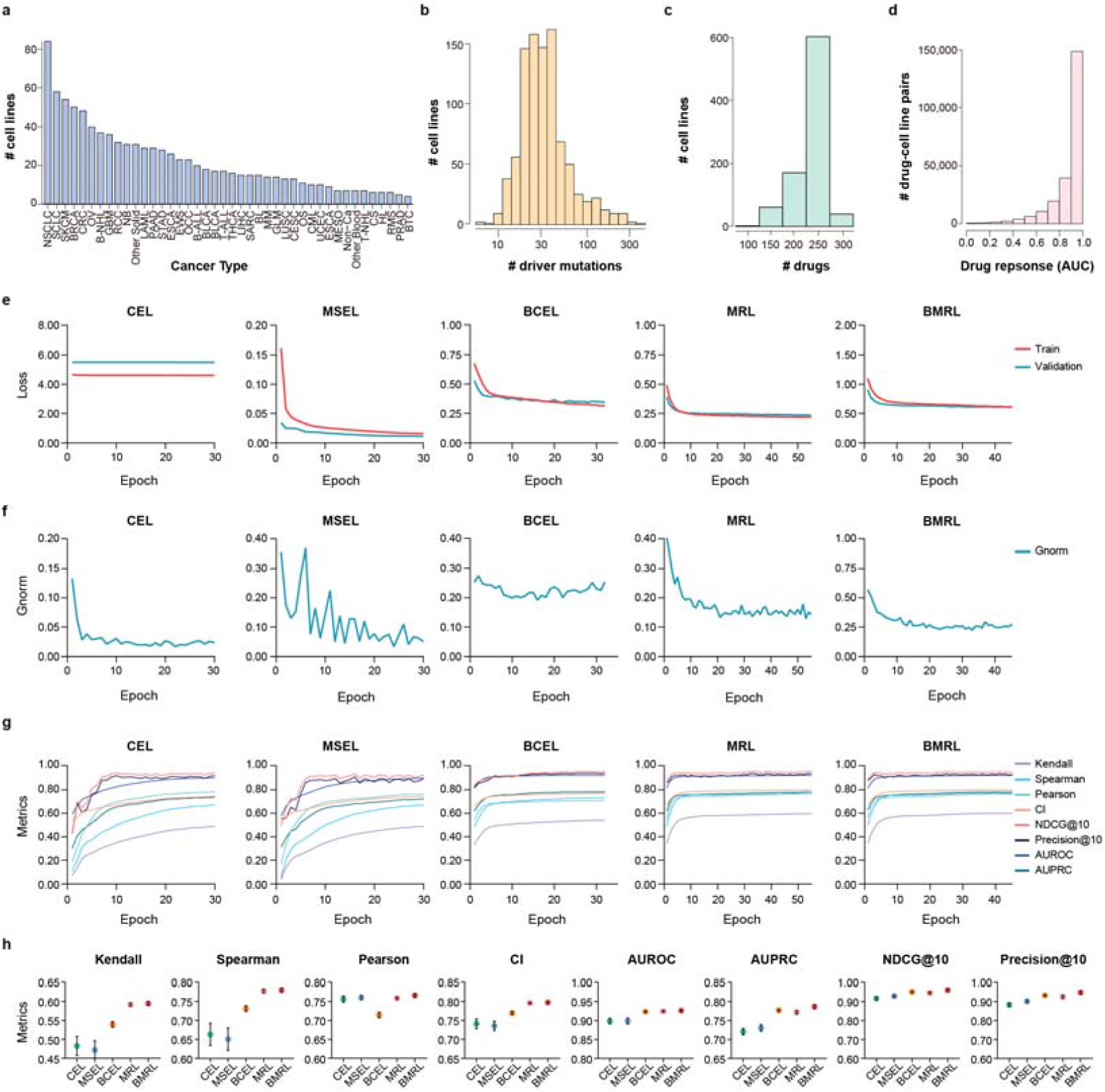
DEPICT training data and optimization performance. **a**, Cancer-type distribution of the CCLE cell lines used for training. **b**, Distribution of the number of driver-gene alterations per cell line. **c**, Distribution of the number of drugs tested per cell line. **d**, Distribution of drug-response AUC values across all cell line-drug pairs. **e**, Training-loss trajectories for the indicated optimization objectives. **f**, Gradient-norm trajectories during training. **g**, Performance metrics across training epochs for the indicated objectives. **h**, Performance of models trained with the indicated loss functions, evaluated by Kendall, Spearman and Pearson correlations, concordance index, AUROC, AUPRC, NDCG@10 and Precision@10.

**Extended Data Fig. 2.**
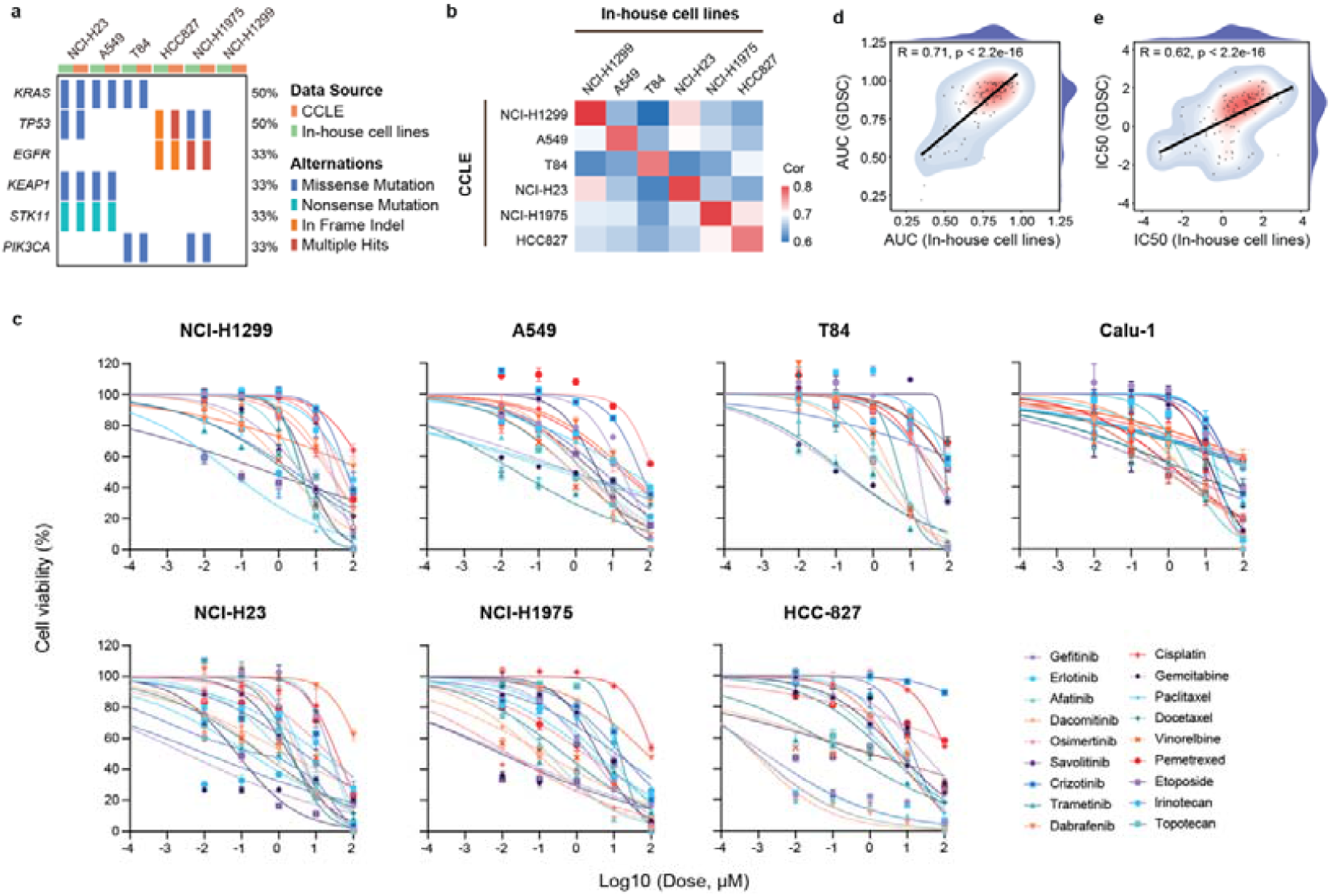
Concordance between in-house datasets and public pharmacogenomic resources. **a**, Key somatic alterations identified by whole-exome sequencing in in-house and matched CCLE cell lines. **b**, Transcriptomic similarity between in-house RNA-seq profiles and CCLE data. **c**, Dose-response curves for 18 drugs across seven lung cancer cell lines. **d, e**, Correlation of in-house and public-database AUC (**d**) and IC50 (**e**) measurements across cell lines and drugs.

**Extended Data Fig. 3.**
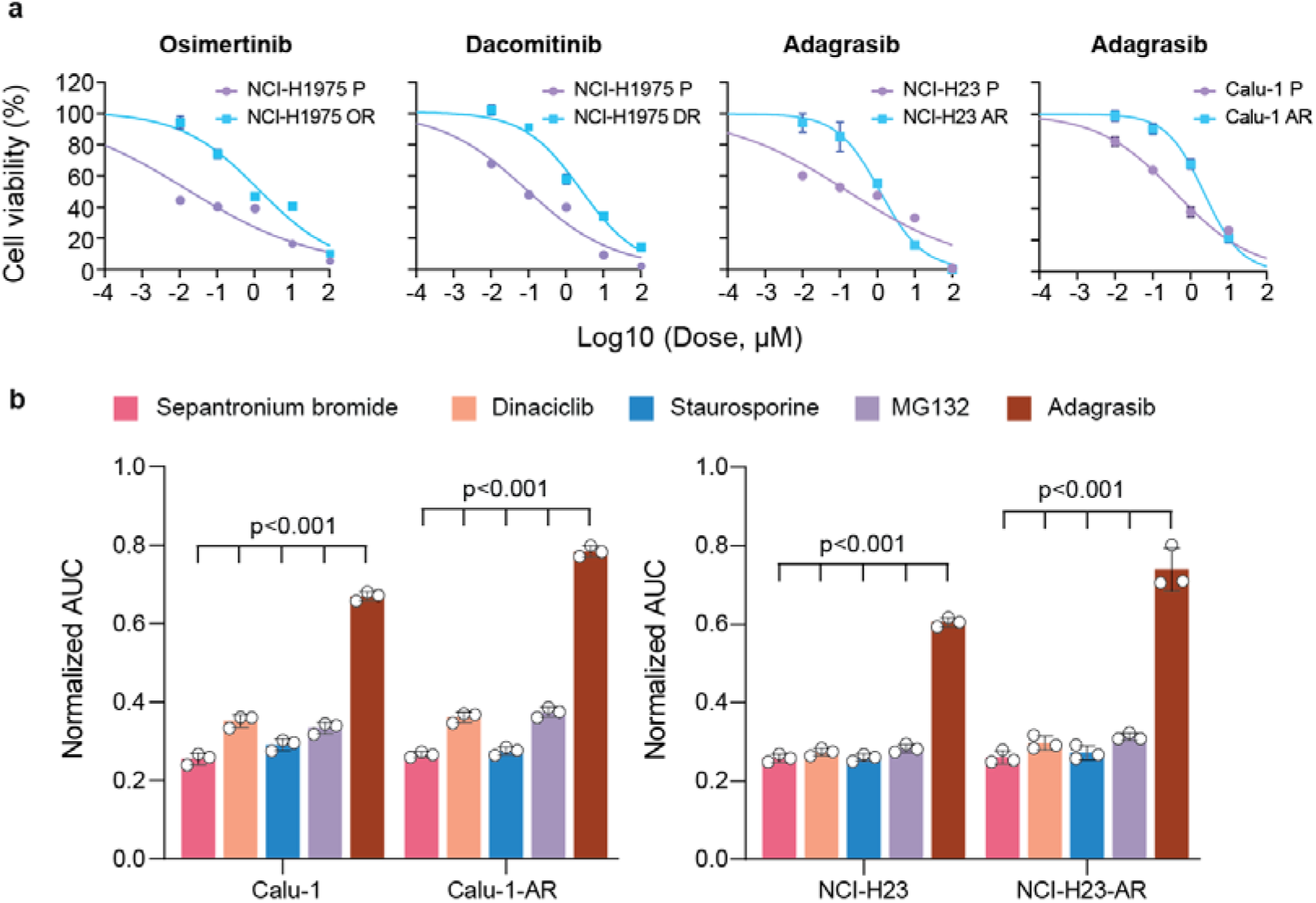
Experimental characterization of acquired resistance and candidate drug responses. **a**, Dose-response curves for parental and resistant NCI-H1975, NCI-H23 and Calu-1 cells treated with their selecting drugs. Cell viability is shown relative to untreated controls. P, parental; OR, osimertinib resistant; DR, dacomitinib resistant; AR, adagrasib resistant. **b**, Responses of parental and adagrasib-resistant Calu-1 and NCI-H23 cells to selected DEPICT-prioritized compounds, shown as normalized AUC. Lower values indicate greater sensitivity. One-way ANOVA.

**Extended Data Fig. 4.**
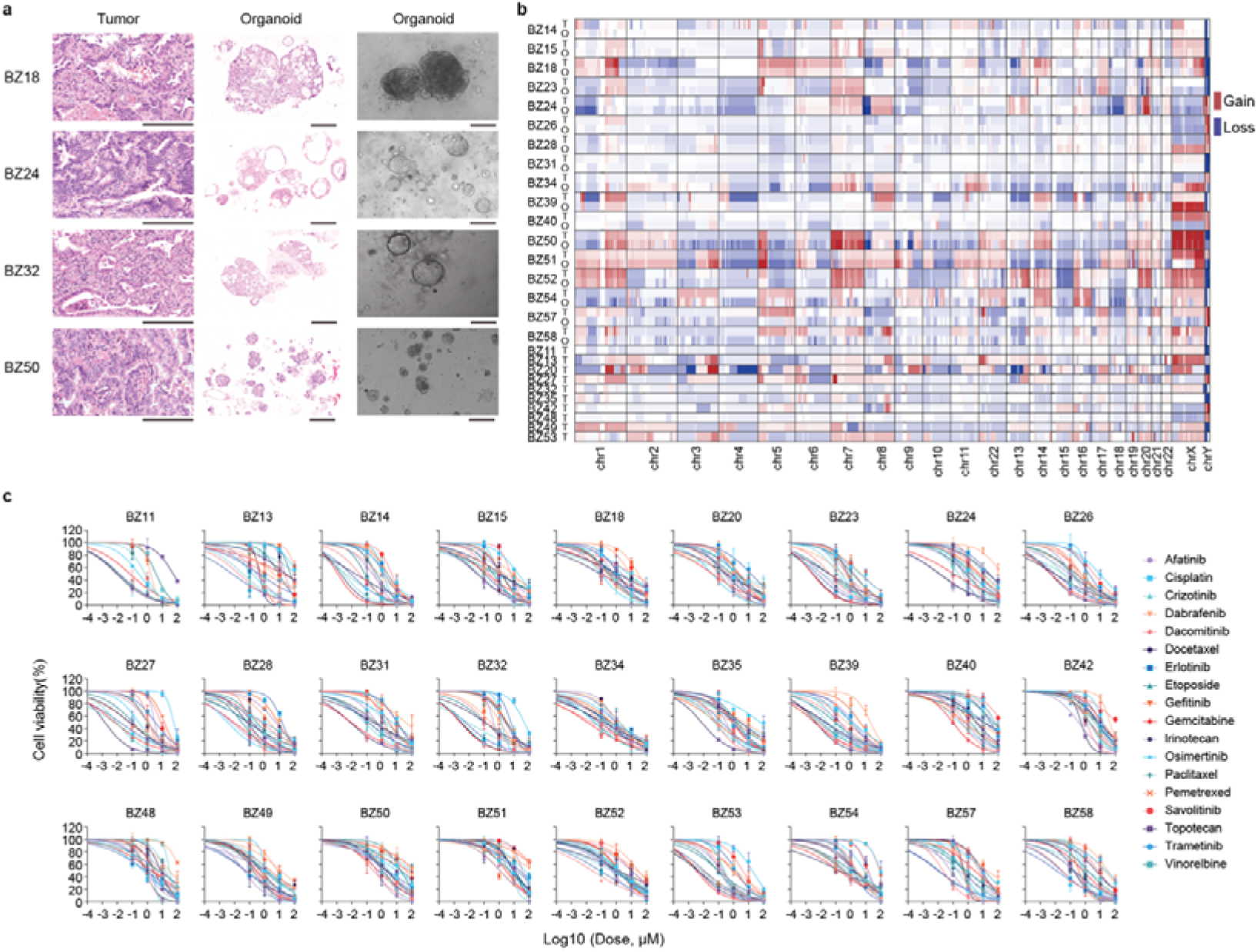
Histopathological and genomic fidelity and drug response profiles of patient-derived lung cancer organoids. **a**, Representative H&E staining of primary tumors and matched PDO sections, together with bright-field images of the corresponding cultures. Scale bars, 100 μm. **b**, Genome-wide copy number profiles of matched tumors (T) and PDOs (O); gains and losses are shown in red and blue, respectively. **c**, Dose-response curves for 18 targeted and chemotherapeutic agents across 27 PDO models.

**Extended Data Fig. 5.**
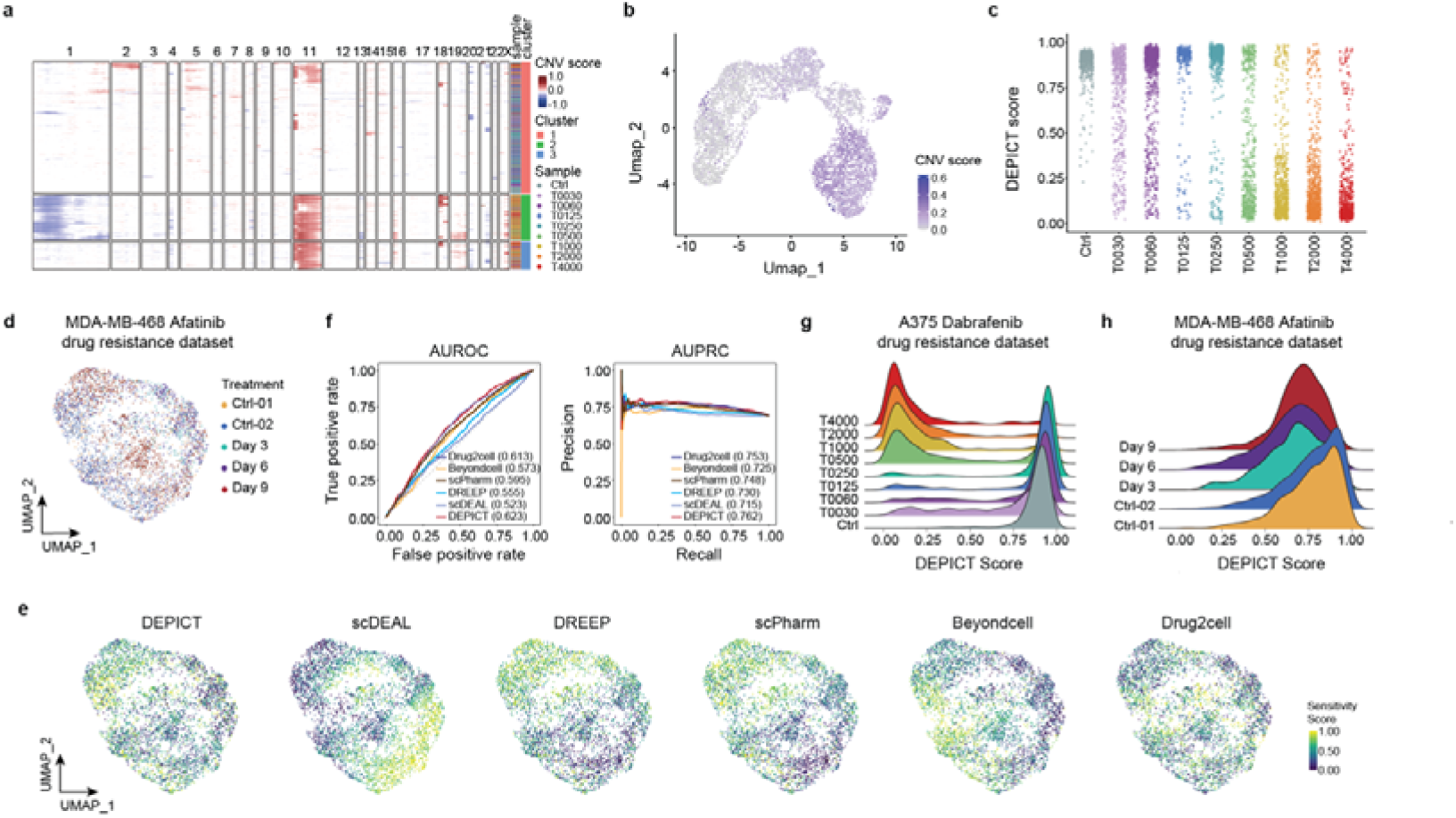
Supporting analyses of single-cell drug sensitivity dynamics in A375 and MDA-MB-468 resistance models. **a**, Copy number variation heatmap of A375 cells exposed to stepwise dabrafenib escalation. **b**, UMAP showing single-cell copy number variation scores in the A375 dataset. **c**, DEPICT-predicted dabrafenib sensitivity scores across A375 treatment conditions. **d**, UMAP of the MDA-MB-468 afatinib resistance dataset across treatment time points. **e**, UMAPs showing sensitivity scores predicted by DEPICT, scDEAL, DREEP, scPharm, Beyondcell and Drug2cell in MDA-MB-468 cells. **f**, AUROC and AUPRC for predicting lineage-tracing-derived survival states in the MDA-MB-468 dataset. **g, h**, Ridge plots of DEPICT-predicted sensitivity across A375 dabrafenib concentrations (**g**) and MDA-MB-468 afatinib treatment time points (**h**).

**Extended Data Fig. 6.**
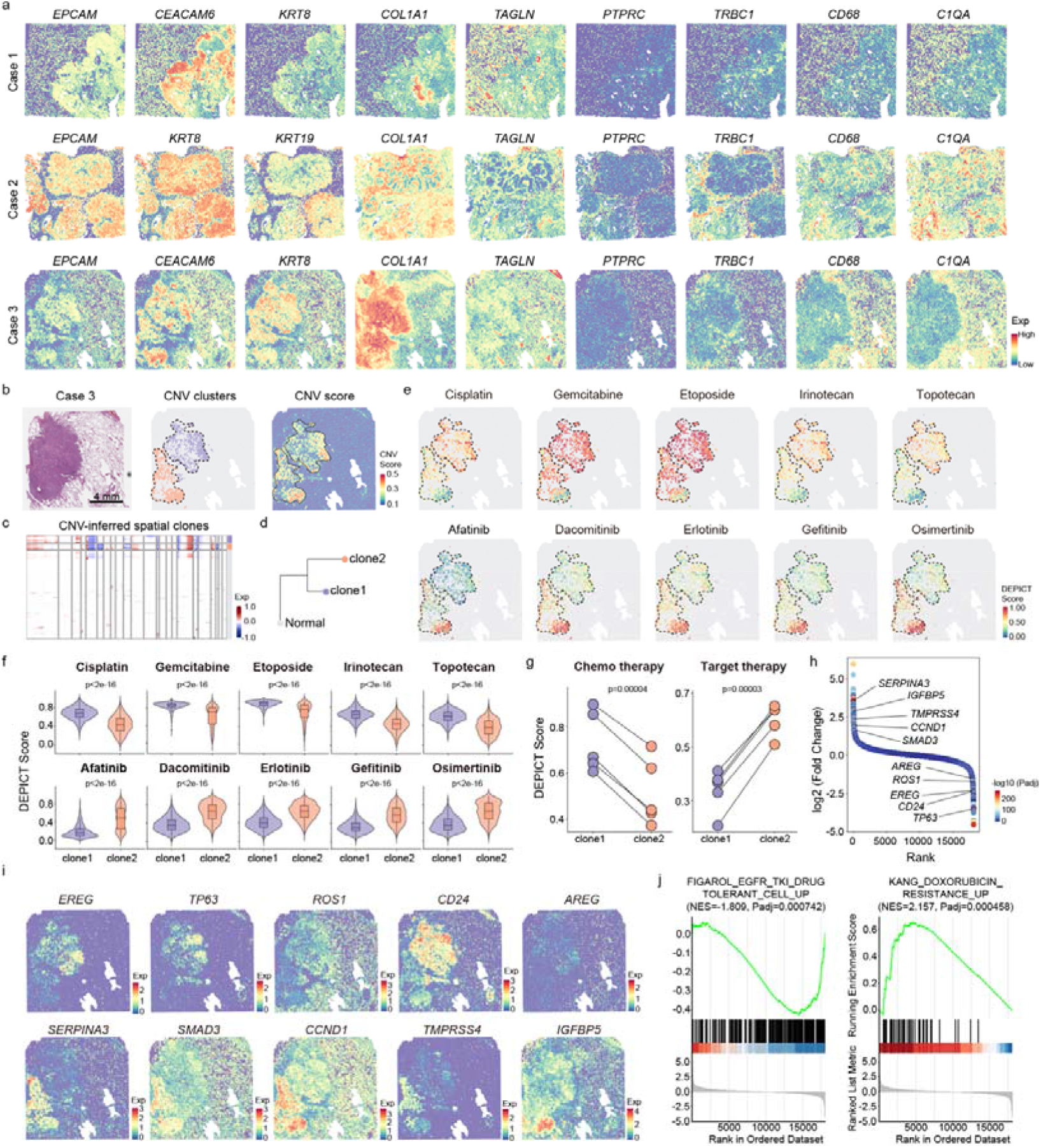
Spatially adjacent tumor clones show divergent predicted responses to chemotherapy and EGFR-targeted therapy. **a**, Spatial expression of representative epithelial, stromal and immune markers across three lung adenocarcinomas. **b**, H&E staining, fastCNV-inferred clone assignment and CNV score for case 3, identifying two adjacent clones. **c**, fastCNV-inferred CNV heatmap. **d**, Phylogenetic tree of normal regions and the two tumor clones. **e**, Predicted sensitivity maps for five chemotherapeutic agents and five EGFR-TKIs. **f**, Predicted drug sensitivity scores in the two clones. Two-sided Wilcoxon rank-sum tests. **g**, Mean chemotherapy and targeted-therapy scores in the two clones. Two-sided paired Student’s t-test. **h**, Differentially expressed genes between the two clones. **i**, Spatial expression of representative clone-associated genes. **j**, Gene set enrichment analysis of clone-associated transcriptional programs.

